# Metabolic free energy and biological codes: a ‘Data Rate Theorem’ aging model

**DOI:** 10.1101/003384

**Authors:** Rodrick Wallace

## Abstract

The living state is cognitive at every scale and level of organization. Since it is possible to associate a broad class of cognitive processes with ‘dual’ information sources, many pathologies can be addressed using statistical models based on the Shannon Coding, the Shannon-McMillan Source Coding, the Rate Distortion, and the Data Rate Theorems, as these impose powerful necessary condition constraints on information generation and exchange, and on system control. Deterministic-but-for-error biological codes do not directly invoke cognition, although they may be essential subcomponents within larger cognitive processes. A formal argument, however, places such codes within a similar framework, with metabolic free energy serving as a ‘control signal’ stabilizing biochemical code-and-translator dynamics in the presence of noise. Demand beyond available energy supply then expresses itself in punctuated destabilization of the coding channel, affecting a spectrum of essential biological functions. Aging, normal or prematurely driven by psychosocial or environmental stressors, must eventually interfere with the routine operation of such mechanisms, triggering chronic diseases associated with senescence. Amyloid fibril formation, intrinsically disordered protein logic gates, and cell surface glycan/lectin ‘kelp bed’ logic gates are reviewed from this perspective. The results, however, generalize beyond coding machineries having easily recognizable symmetry modes, and strip a full layer of mathematical complication from the study of phase transitions in nonequilibrium biological systems.

## 1 Introduction

The understanding of aging has recently become closely intertwined with the understanding of cellular mitochondrial function (D.C. Wallace 2005, 2010). To paraphrase Lee and Wei (2012), aging is a degenerative process that is associated with progressive accumulation of deleterious changes with time, reduction of physiological function and increase in the chance of disease and death. Studies reveal a wide spectrum of alterations in mitochondria and mitochondrial DNA with aging. Mitochondria are the main cellular energy sources that generate the cellular energy source ATP through respiration and oxidative phosphorylation in the inner membrane of mitochondria. The respiratory chain of that system is also the primary intracellular source of reactive oxygen species and free radicals under normal physiological and pathological conditions. In addition, mitochondria play a central role in a great variety of cellular processes.

Numerous biochemical studies on isolated mitochondria revealed that the electron transport activities of respiratory enzyme complexes gradually decline with age in the brain, skeletal muscle, liver and skin fibroblasts of normal human subjects. Numerous molecular studies demonstrated that somatic mutations in mitochondrial DNA accumulate with age in a variety of tissues in humans. These age-associated changes in mitochondria are well correlated with the deteriorative processes of tissues in aging.

However, although abundant experimental data have been gathered in the past decade to support the concept that decline in mitochondrial energy metabolism, reactive oxygen species overproduction and accumulation of mitochondrial DNA mutations in tissue cells are important contributors to human aging, the detailed mechanisms by which these biochemical events cause aging have remained to be established.

Wallace (2014d) applies necessary conditions statistical models from communications and control theory to examine the central role of metabolic free energy (MFE) in biorgulation, using a very general Rate Distortion Theorem development. Here, we delve into details, studying the operation and regulation of a spectrum of biological codes at and across the cellular level of organization. In essence, failure to provide adequate levels of MFE can trigger collapse of critical code-and-translation mechanisms, either to dysfunctional simplified ‘ground state’ modes having collapsed symmetry states, or to highly pathological unstable dynamics.

As Ge and Quan (2011) put it, one of the challenging questions in biological physics is whether nonequilibrium phase transitions play an important role in living systems. We will show this to be very broadly true, with the asymptotic limit theorems of information and control theories stripping away a layer of mathematical complication for the systems to which they apply.

## 2 Cognition and codes

It has long been maintained that the living state is cognitive at every scale and level of organization (e.g., Maturana and Varela 1980; Atlan and Cohen 1998; Wallace 2012c, 2014a). Since it is possible to associate a broad class of cognitive processes with ‘dual’ information sources (e.g., Wallace 2005, 2007, 2012c, 2014a), many phenomena – most particularly, complex patterns of behavioral pathology – can be addressed using statistical models based on the Shannon Coding, the Shannon-McMillan Source Coding, the Rate Distortion, and the Data Rate Theorems (S/SM/RD/DR), as these provide powerful necessary conditions on all information generation and exchange, and on the role of information in system control (e.g., Wallace 2012c, 2014b, c).

Strictly speaking, biological codes, although they may be studied using information theoretic methods, as Tlusty (2007, 2008) does for the genetic code, do not actively invoke cognition, although, as with cognitive gene expression (e.g., Wallace and Wallace 2010), they may become essential subcomponents within larger cognitive processes. Nonetheless, as we will show here, something similar to the Data Rate Theorem that connects information and control theories still applies to biological coding processes, but based on the flow of metabolic free energy rather than on the flow of control information, as for the DRT. See the Mathematical Appendix for an explicit statement of that theorem.

Tlusty’s (2007) information theoretic topological analysis of the genetic code relies on minimizing certain characteristic error measures. Wallace (2012a) examined the role of metabolic free energy in the evolution of such codes, using similar methods. Here we first generalize the argument, based on a Black-Scholes ‘cost’ analysis. We then explore a model of punctuated code failure under free energy constraint that is roughly analogous to Data Rate Theorem (DRT) limitations in control theory (e.g., Nair et al. 2007). This will suggest a deeper understanding of the onset of the chronic diseases of aging, and of those driven by psychosocial or environmental stresses that cause premature aging.

The essential point of the DRT is the unification of control and information theories, finding that certain kinds of unstable systems cannot be stabilized if the rate of control information is below a critical limit, defined as the ‘topological information’ generated by the unstable system. Metabolic free energy plays a surprisingly similar role in stabilizing deterministic-but-for-error biological codes.

Although we will focus primarily on codes having a relatively easily characterized symmetry structure, the general argument should apply as well to far less symmetric codes, in the sense of Tomkins (1975).

Tlusty’s (2007) central idea is that

> To discuss the topology of errors we portray the codon space as a graph whose verticies are the codons… Two codons… are linked by an edge if they are likely to be confused by misreading… We assume that two codons are most likely to be confused if all their letters except for one agree and therefore draw an edge between them. The resulting graph is natural for considering the impact of translation errors on mutations because such errors almost always involve a single letter difference, that is, a movement along an edge of the graph to a neighboring vertex.
>
> The topology of a graph is characterized by its genus *γ*, the minimal number of holes required for a surface to embed the graph such that no two edges cross. The more connected that a graph is the more holes are required for its minimal embedding… [T]he highly interconnected 64-codon graph is embedded in a holey, *γ* = 41 surface. The genus is somewhat reduced to *γ* = 25 if we consider only 48 effective codons…
>
> The maximum [of an information-theoretic functional] determines a single contiguous domain where a certain amino acid is encoded… Thus every mode corresponds to an amino acid and the number of modes is the number of amino acids. This compact organization is advantageous because misreading of one codon as another codon within the same domain has no deleterious impact. For example, if the code has two amino acids, it is evident that the error-load of an arrangement where there are two large contiguous regions, each coding for a different amino acid, is much smaller than a ‘checkerboard’ arrangement of the amino acids.

This is analogous to the well-known topological coloring problem. However, in the coding problem one desires maximal similarity in the colors of neighboring ‘countries’, while in the coloring problem one must color neighboring countries by different colors. After some development (Tlusty 2008), the number of possible amino acids in this scheme is determined by Heawood’s formula (Ringel and Young 1968). Explicitly,

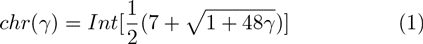

where *chr*(*γ*) is the number of ‘colored’ regions, *Int* is the integer value of the enclosed expression, and *γ* is the genus of the surface, roughly speaking, the number of ‘holes’. In general, *γ* =1 − (1/2)(*V* − *E* + *F*), where *V* is the number of code network vertices, *E* the number of network edges, and *F* the number of enclosed faces.

Tlusty (2007) models the emergence of the genetic code as a transition in a noisy information channel, using an approach based on the Rate Distortion Theorem, with the optimal code is described by the minimum of a ‘free energy’-like functional, allowing description of the code’s emergence as a transition akin to a phase transition in statistical physics. The basis for this is the observation that a supercritical phase transition is known to take place in noisy information channels.

The noisy channel is controlled by a temperature-like parameter that determines the balance between the information rate and the distortion in the same way that physical temperature controls the balance between energy and entropy in a physical system. Following Tlusty’s equation (2), the free energy functional has the form *D* – *TS* where *D* is the average error load’, equivalent to average distortion in a rate distortion problem, *S* is the ‘entropy due to random drift’, and *T* measures the strength of random drift relative to the selection force that pushes towards fitness maximization. This is essentially a Morse function (Pettini 2007; Matsumoto 2002). According to Tlusty’s analysis, at high *T* the channel is totally random and it conveys zero information. At a certain critical temperature *T_c_* the information rate starts to increase continuously.

The average distortion *D* measures the average difference between the genetic ‘message’ sent by a complicated codon ‘statement’ and what is actually expressed by the genetic (and epigenetic) translation machinery in terms of an amino acid sequence. See figure 1.

**Figure 1:**
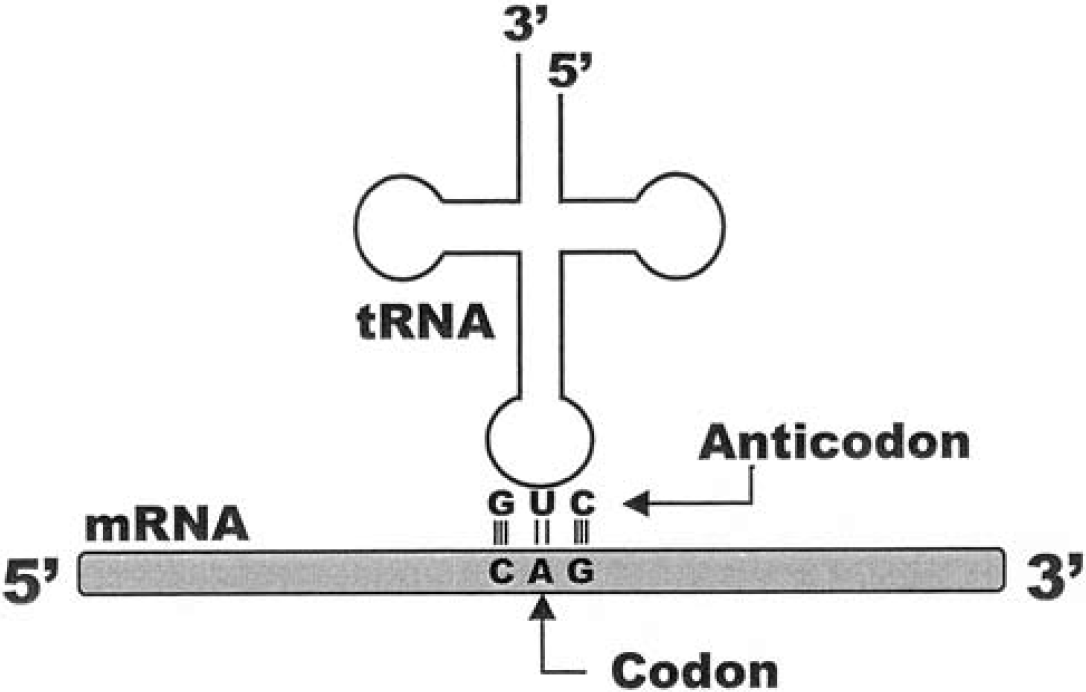
Adapted from fig. 1.8 of Shmulevich and Dougherty (2007). DNA meets RNA in modern protein synthesis. The anticodon at one end of a tRNA molecule binds to its complementary codon in mRNA derived directly from the genome. The average distortion *D* is a measure of the difference between what is supposed to be coded by a genome sequence and what is actually expressed as an amino acid sequence. Sequence-to-sequence translation is not highly parallel, in this model, and the process can be well characterized by the rate distortion function *R*(*D*) representing the minimum channel capacity needed to produce average distortion less than *D.* Similar, but more complex, biological matching occurs with intrinsically disordered protein reactions, and at the cell surface, with glycan/lectin interaction. It is, however, possible to imagine translation-at-a-distance in which molecular species do not interact directly, but via intermediate chemical signals, in the sense of Tomkins (1975).

Here we envision a multi-step process in which the rate distortion function *R*(*D*) – described more fully in the next section – between codon sequence and amino acid sequence plays the central role. In the first step, *R*(*D*), a nominally extensive quantity, but one physically limited by the channel construction of figure 1, serves as a temperature-analog in a one-parameter distribution of information source uncertainties representing different coding strategies, from which a free energy functional is constructed. While *R*(*D*) is not ‘temperature-like’ – e.g., under a given circumstance it can be increased as much as one likes by establishing parallel channels – the physical structure of translation constrains that approach, ensuring the ‘locally intensive’ nature of the rate distortion function. Pettini’s (2007) ‘topological hypothesis’ implies that topological shifts in code structure accompany phase transitions in a free energy functional constructed from the distribution of information source uncertainties arising from possible code topologies.

The second stage of the argument revolves around the relation between intensive indices of metabolic free energy availability – e.g., underlying energy per molecular transaction, and/or efficiency of its use – and *R*(*D*), leading to a second free energy-like functional that undergoes another set of punctuated phase changes.

While the genetic code has received much attention, Hecht et al. (2004) note that protein α-helices have the ‘code’ 101100100110… where 1 indicates a polar and 0 a nonpolar amino acid. Protein *β*-sheets have the simpler coding 10101010… Wallace (2010), in fact, extends Tlusty’s topological analysis via Heawood’s graph genus formula to the more complicated protein folding classifications of Chou and Maggiora (1998). Wallace (2012b) argues, in addition, that a similar argument must apply to the basic monosaccharides associated with glycan ‘kelp frond’ production at the surface of the cell. Wallace (2012d), as we shall show, provides machinery for including intrinsically disordered protein and glycan/lectin cell surface reactions within the overall perspective. Again, here we shall be interested in calculating metabolic costs necessarily associated with limiting error across such biological codes, and will model both punctuated code evolution and a form of instability triggered by metabolic energy restriction, or by the growth of energy demand beyond available resources.

## 3 Some information theory

The existence of a code implies the existence of an information source using that code, and the behavior of such sources is constrained by the asymptotic limit theorems of information theory. That is, the interaction between biological subsystems associated with a code can be formally restated in communication theory terms. Essentially, observation of a code directly implies existence of an information source using it.

Think of the machinery listing a sequence of codons as communicating with machinery that produces amino acids, and compare what is actually produced with what should have been produced, perhaps by a simple survival of the fittest selection mechanism, perhaps via some more sophisticated error-correcting systems.

Suppose a sequence of signals is generated by a biological information source *Y* having output *y*^*n*^ = *y_1_, y_2_*, … – codons. This is ‘digitized’ in terms of the observed behavior of the system with which it communicates, for example a sequence of ‘observed behaviors’ *b*^*n*^ = *b_1_, b_2_*, … – amino acids. Assume each *b*^*n*^ is then deterministically retranslated back into a reproduction of the original biological signal, 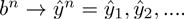

Define a distortion measure 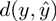 which compares the original to the retranslated path. Many distortion measures are possible. The Hamming distortion is defined simply as

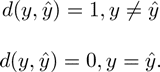

For continuous variates the squared error distortion is just 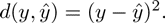

There are many possible distortion measures. The distortion between paths *y*^*n*^ and 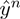 is defined as 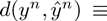 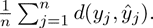

A remarkable characteristic of the Rate Distortion Theorem is that the basic result is independent of the exact distortion measure chosen (Cover and Thomas 2006).

Suppose that with each path *y*^*n*^ and *b*^*n*^-path retranslation into the *y*-language, denoted 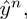 there are associated individual, joint, and conditional probability distributions 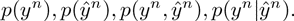.

The average distortion is defined as

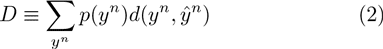

This is essentially the ‘error load’ of Tlusty’s (2007) equation (1).

It is possible to define the information transmitted from the *Y* to the 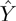 process using the Shannon source uncertainty of the strings:

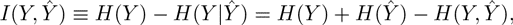

where *H*(…, …) is the standard joint, and *H*(…|…) the conditional, Shannon uncertainties (Cover and Thomas 2006).

If there is no uncertainty in *Y* given the retranslation 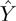, then no information is lost, and the systems are in perfect synchrony.

In general, of course, this will not be true.

The rate distortion function *R*(*D*) for a source *Y* with a distortion measure 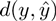 is defined as

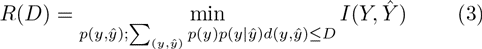

The minimization is over all conditional distributions 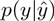 for which the joint distribution 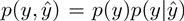 satisfies the average distortion constraint (i.e., average distortion *≤ D*).

The Rate Distortion Theorem states that *R*(*D*) is the minimum necessary rate of information transmission which ensures the communication between the biological vesicles does not exceed average distortion *D*. Thus *R*(*D*) defines a minimum necessary channel capacity. Cover and Thomas (2006) or Dembo and Zeitouni (1998) provide details. The rate distortion function has been calculated for a number of systems.

Cover and Thomas (2006, Lemma 13.4.1) show that *R*(*D*) is necessarily a decreasing convex function of *D* for any reasonable definition of distortion.

That is, *R*(*D*) is always a reverse J-shaped curve. This will prove crucial for the overall argument. Indeed, convexity is an exceedingly powerful mathematical condition, and permits deep inference (e.g., Rockafellar 1970). Ellis (1985, Ch. VI) applies convexity theory to conventional statistical mechanics.

For a Gaussian channel having noise with zero mean and variance *σ*^2^, using the squared distortion measure,

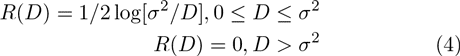

Recall the relation between information source uncertainty and channel capacity (Cover and Thomas 2006):

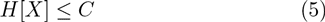

where *H* is the uncertainty of the source *X* and *C* the channel capacity. Remember also that

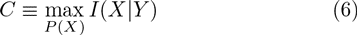

where *P*(*X*) is chosen so as to maximize the rate of information transmission along a channel *Y*.

Note that for a parallel set of noninteracting channels, the overall channel capacity is the sum of the individual capacities, providing a ‘consensus average’ that does not apply in the case of modern molecular coding.

Finally, recall the analogous definition of the rate distortion function above, again an extremum over a probability distribution.

Recall also the homology between information source uncertainty and free energy density. More formally, if *N*(*n*) is the number of high probability ‘meaningful’ – that is, grammatical and syntactical – sequences of length *n* emitted by an information source *X*, then, according to the Shannon-McMillan Theorem, the zero-error limit of the Rate Distortion Theorem,

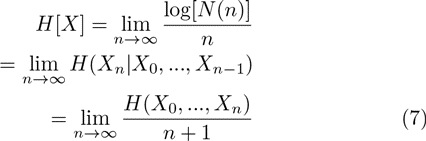

where*H*(…|…) is the conditional and *H*(…, …) is the joint Shannon uncertainty.

In the limit of large *n*, *H*[*X*] becomes homologous to the free energy density of a physical system at the thermodynamic limit of infinite volume. More explicitly, the free energy density of a physical system having volume *V* and partition function *Ζ*(*β*) derived from the system’s Hamiltonian – the energy function – at inverse temperature *β* is (e.g., Pettini, 2007)

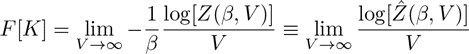

with 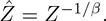 The latter expression is formally similar to the first part of equation (7), a matter having deep implications: Feynman (2000) describes in great detail how information and free energy have an inherent duality. Feynman, in fact, defines information precisely as the free energy needed to erase a message. The argument is surprisingly direct, and for very simple systems it is easy to design a small (idealized) machine that turns the information within a message directly into usable work – free energy. Information is a form of free energy and the construction and transmission of information within living things consumes metabolic free energy, with nearly inevitable losses via the second law of thermodynamics. If there are limits on available metabolic free energy there will necessarily be limits on the ability of living things to process information.

From one perspective, the Shannon-McMillan Theorem can be said to define a nonequilibrium steady state that can undergo nonequilibrium phase transitions in the sense of Ge and Qian (2011). These are analogous to, but different from those of physical systems. A similar development produces an analog to Onsager-like nonequilibrium theomodynamic relations.

## 4 Groupoid symmetry shifting

Here we follow, in part, the argument of Wallace (2012a). The direct model finds codons generated by a black box information source whose source uncertainty is constrained by the richness of the coding scheme of Tlusty’s analysis. More complex codes will be associated with higher information source uncertainties, i.e., the ability to ‘say’ more in less time, using a more complicated coding scheme. Suppose there are *n* possible coding schemes. The simplest approach is to assume that, for a given rate distortion function and distortion measure, *R*(*D*), under the constraints of figure 1, serves much as an external temperature bath for the possible distribution of information sources, the set *{H_1_, …, H_n_}.* That is, low distortion, represented by a high rate of transmission of information between codon machine and amino acid machine, permits more complicated coding schemes according to a classic Gibbs relation

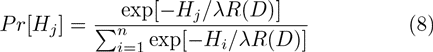

where *Pr*[*H*_*j*_] is the probability of coding scheme *j* having information source uncertainty *H*_*j*_.

We assume that *Pr*[*H*_*j*_] is a one parameter distribution in the ‘extensive’ quantity *R*(*D*) (monotonic convex, however, in *D*) rather than a simple ‘intensive’ temperature-analog. This is permitted under the ‘structurally intensive’ circumstance of figure 1.

The free energy-like Morse Function *F*_*R*_ associated with this probability is defined as

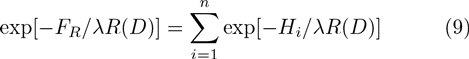

Applying Landau’s spontaneous symmetry lifting argument to *F*_*R*_ (Pettini 2007) generates topological transitions in codon graph structure as the ‘temperature’ *R*(*D*) increases, i.e., as the average distortion *D* declines, via the inherent convexity of the Rate Distortion Function. That is, as the channel capacity connecting codon machines with amino acid machines increases, more complex coding schemes become possible:

1. The genus of the embedding surface for a topological code can be expressed in terms of the Euler characteristic of the manifold, 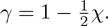
2. χ can be expressed in terms of the cohomology structure of the manifold (Lee 2000, Theorem 13.38).
3. By the Poincare Duality Theorem, the homology groups of a manifold are related to the cohomology groups in the complementary dimension (Bredon 1993, p. 348).
4. The (co)homology groupoid can be taken as the disjoint union of the (co)homology groups of the embedding manifold.

One can then invert Landau’s Spontaneous Symmetry Breaking arguments and apply them to the (co)homology groupoid in terms of the rising ‘temperature’ *R*(*D*), to obtain a punctuated shift to increasingly complex genetic codes with increasing channel capacity. See the Mathematical Appendix for a summary of standard material on groupoids. Brown (1987) and Weinstein (1996) provide more detail.

What, then, drives *R*(*D*), as this, in turn, drives punctuated changes in the genetic code? Here we will significantly diverge from the arguments in Wallace (2012a), invoking a Black-Scholes formalism for ‘cost’ in terms of demand for metabolic free energy. Later, we will use a similar argument to examine failures in the dynamics of evolutionarily fixed codes under free energy restraints.

## 5 Metabolic energy costs

Suppose that metabolic free energy is available at a rate 𝓗 determined by environmental structure and previous evolutionary trajectory, which may be prior to the emergence of efficient photosynthesis, predation, mutualism, parasitism, and the like. We iterate the treatment and consider 𝓗 as the temperature analog in a Landau model on the Rate Distortion Function itself. That is, let *R*(*D*) be the Rate Distortion Function describing the relation between system intent and system impact. This is essentially a channel capacity, and information transmission rate between the coding machine and the structure or structures that biological code is attempting to affect.

The distortion represents the dynamics of the disjunction between the intent of a code and the actual productions of the system. Let *R*_*t*_ be the RDF of the channel connecting them at time *t*. The relation can, under conditions of both white noise and volatility, be expressed as

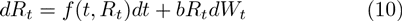

Let *𝓗(*R*_*t*_,t*) represent the rate of incoming metabolic free energy that is needed to achieve *R*_*t*_ at time *t*, and expand using the Ito chain rule:

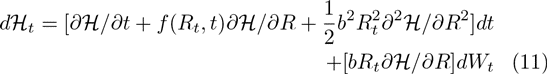

Define *𝓛* as the Legendre transform of the free energy rate 𝓗, a kind of entropy, having the form

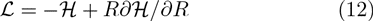

Using the heuristic of replacing *dX* with Δ*Χ* in these expressions, and applying the results of equation (11), produces:

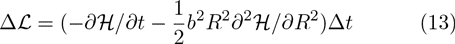

Analogous to the Black-Scholes calculation, the terms in *f* and *dW_t_* cancel out, so that the effects of noise are subsumed in the Ito correction involving *b*. Clearly, however, this also invokes powerful regularity assumptions that may often be violated. Matters then revolve about model robustness in the face of such violation.

*𝓛*, as the Legendre transform of the free energy rate measure 𝓗, is a kind of entropy that can be expected to rapidly reach an extremum at nonequilibrium steady state (nss). There, Δ*L/Δt* = *∂𝓗/∂t* = 0, so that

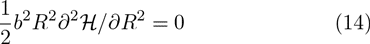

having the solution

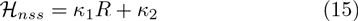

This ‘simple’ result, we will show, permits a Landau-analog phase transition analysis in which the metabolic free energy available from the embedding ecosystem serves to raise or lower the possible richness of a system’s possible biological codes. As Wallace (2012a) argues, if 𝓗 is relatively large then there are many possible complex codes. If, however, sufficient metabolic free energy is not available, then the system can only entertain a few simplified codings.

While the aerobic transition apparently enabled endosymbiotic processes producing eukaryotic organisms, it may also have enabled evolution of the extraordinarily rich glycan/lectin cell surface codings essential to all multicellular organisms. Wallace (2012b), however, infers a paradox: that full coding, having 5,000-7,500 ‘glycan determinant’ amino acid analogs made up of the appropriate basic set of monosaccharides, would require a coding manifold with topological genus in the millions, suggesting the need for an intermediate layer of cognitive mechanism at the cell surface.

## 6 Code/translator stability

Van den Broeck et al. (1994, 1997), Horsthemeke and Lefever (2006), and others, have noted that analogous results relating phase transition to driving parameters in physical systems can be obtained by using the rich stability criteria of stochastic differential equations.

The motivation for this approach is the observation that a Gaussian channel with noise variance *σ*^2^ and zero mean has a Rate Distortion function *R*(*D*) = 1/2log[*σ*^2^/*D*] using the squared distortion measure for the average distortion *D*. Defining a ‘Rate Distortion entropy’ as the Legendre transform

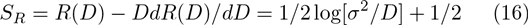

the simplest possible nonequilibrium Onsager equation (de Groot and Mazur 1984) is just

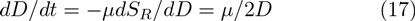

where *t* is the time and *μ* is a diffusion coefficient. By inspection, 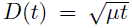, the classic solution to the diffusion equation. Such ‘correspondence reduction’ serves as a base to argue upward in both scale and complexity.

But deterministic coding does not involve diffusive drift of average distortion. Let 𝓗 again be the rate of available metabolic free energy. Then a plausible model, in the presence of an internal system noise *β*^2^ in addition to the environmental channel noise defined by *σ*^2^, is the stochastic differential equation

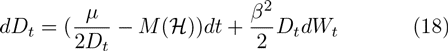

where *dW_t_* represents unstructured white noise and *M*(𝓗) ≥ 0 is monotonically increasing.

This has the nonequilibrium steady state expectation

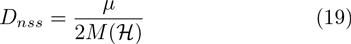

Using the Ito chain rule on equation (18) (Protter 1990; Khasminskii 2012), one can calculate the variance in the distortion as 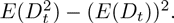

Letting 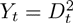 and applying the Ito relation,

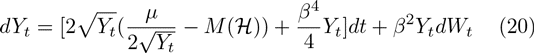

where (*β*^4^/4)*Y_t_* is the Ito correction to the time term of the SDE.

A little algebra shows that no real number solution for the expectation of 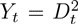 can exist unless the discriminant of the resulting quadratic equation is ≥ 0, producing a minimum necessary rate of available metabolic free energy for stability defined by

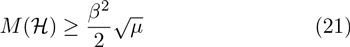

Values of *M*(𝓗) below this limit will trigger a phase transition into a less integrated – or at least behaviorally different – system in a highly punctuated manner, much as in the Landau example.

From equations (15) and (19),

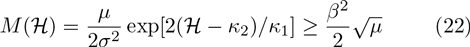

Solving for 𝓗 gives the necessary condition

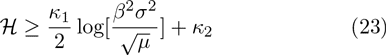

for there to be a real second moment in *D*, under the subsidiary condition that 𝓗 ≥ 0.

Given the context of this analysis, failure to provide adequate rates of metabolic free energy 𝓗 would represent the onset of a regulatory stability catastrophe. The corollary, of course, is that environmental influences increasing *β*, *σ*, the *κ*_*i*_, or reducing *μ*, would be expected to overwhelm internal controls, triggering similar instability.

Variations of the model are possible, for example, applying the formalism to the ‘natural’ channel, having the rate distortion function *R*(*D*) = *σ*^2^/*D*. The calculation is direct.

Equation (23) is a close analog to the Data Rate Theorem (Nair et al. 2007, Theorem 1). Again, see the Mathematical Appendix for a statement of that theorem. The implication is that there is a critical rate of available metabolic free energy below which there does not exist any quantization, coding, or control scheme, able to stabilize an (inherently) unstable biological system.

Normal, or stress-induced, aging would, at some point, be expected to affect the magnitudes of the parameters on the right hand side of the expression in equation (23), while simultaneously decreasing the ability to provide metabolic free energy – decreasing 𝓗. This would result in onset of serious dysfunctions across a range of scales and levels of organization, from genetic to protein folding to cell surface signaling, and beyond (Tomkins 1975).

## 7 Extending the model

It is possible to reinterpret the results of equation (23) from the perspective of Section 3, producing a more general picture of code failure under metabolic energy limitations. Suppose we agree that equation (15) is only a first approximation, and that we can take the Rate Distortion Function *R* as a monotonic increasing function of available metabolic free energy rate 𝓗 that we begin to interpret as an effective system ‘temperature’. Suppose also there are very many more possible ‘modes’ of code behavior, in addition to the simple stability/instability break point implied by equation (23). That is, we now expect complex ‘phase transitions’ in code function with either changing demand for, or ability to provide, rates of metabolic free energy to the coding/translating machine(s).

Given a large enough set of possible modes of code/translation behavior, we write another pseudoprobability like equation (8),

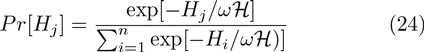

where *H_j_* is the source uncertainty to be associated with each functional mode *j*.

This leads to another ‘free energy’ Morse Function, 𝓕, defined now in terms of the rate of available metabolic free energy

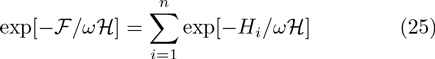

Certain details of information phase transitions for this system can be calculated using ‘biological’ renormalization methods (Wallace, 2005, Section 4.2) analogous to, but much different from, those used in the determination of physical phase transition universality classes (Wilson, 1971).

Given *𝓕* as a free energy analog, what are the transitions between functional realms? Suppose, in classic manner, it is possible to define a characteristic ‘length’, say *l*, on the system. It is then possible to define renormalization symmetries in terms of the ‘clumping’ transformation, so that, for clumps of size *L*, in an external ‘field’ of strength *J* (that can be set to 0 in the limit), one can write, in the usual manner (e.g., Wilson 1971)

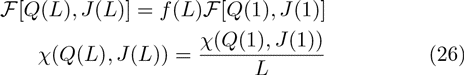

where χ is a characteristic correlation length and *Q* is an ‘inverse temperature measure’, i.e., ∝ 1/*ω𝓗*.

As described in Wallace (2005), very many ‘biological’ renormalizations, *f*(*L*), are possible that lead to a number of quite different universality classes for phase transition. Indeed, a ‘universality class tuning’ can be used as a tool for large-scale regulation of the system. While Wilson (1971) necessarily uses *f*(*L*) ∝ *L*^3^ for simple physical systems, following Wallace (2005), it is possible to argue that, since *𝓕* is so closely related to information measures, it is likely to ‘top out’ at different rates with increasing system size, so other forms of *f*(*L*) must be explored. Indeed, standard renormalization calculations for *f*(*L*) ∝ *L*^*δ*^, *m* log(*L*) + 1, and exp[*m*(*L* − 1)/*L*] all carry through.

This line of argument leads to complex forms of highly punctuated phase transition in code/translator function with changes in demand for, or supply of, the metabolic free energy needed to run the machine.

## 8 Amyloid fibril formation

Another possible inference from the considerations of Sections 3 and 6 is that, under MFE inadequacy, grossly simplified ‘defacto’ codes may sometimes begin to operate in place of the full code. The most direct example, perhaps, is the collapse of the ‘protein folding code’ from the relatively complicated symmetries described in Wallace (2010) to *β*-sheet amyloid plaques and fibrils in many protein folding disorders.

More specifically, globular proteins, following the observations of Chou and Maggiora (1998), have four major, and perhaps as many as another six minor, classifications. This suggests a Tlusty code error network that is, essentially, a large ‘sphere’, having one minor, and possibly as many as three more subminor attachment handles, according to Heawood’s formula. These basic structures build a highly complicated ‘protein world’ that cannot be simply characterized.

The prebiotic ‘amyloid world’ of Maury (2009), in contrast, is built on a single *β*-sheet structure, and shows, by contrast to the protein world, in its full extent, a simple eight-fold steric zipper (Sawaya et al. 2007).

As Goldschmidt et al. (2010) put the matter,

> We found that [protein segments with high fibrillation propensity] tend to be buried or twisted into unfavorable conformations for forming beta sheets… For some proteins a delicate balance between protein folding and misfolding exists that can be tipped by changes in environment, destabilizing mutations, or even protein concentration…
>
> In addition to the self-chaperoning effects described above, proteins are also protected from fibrillation during the process of folding by molecular chaperones…
>
> Our genome-wide analysis revealed that self-complementary segments are found in almost all proteins, yet not all proteins are amyloids. The implication is that chaperoning effects have evolved to constrain self-complementary segments from interaction with each other.

Clearly, effective chaperoning requires considerable metabolic energy, and failure to provide levels adequate for both maintaining and operating such biochemical translation machinery would trigger a canonical ‘code collapse’, most likely in a highly punctuated manner with available metabolic free energy.

Indeed, Budrikis et al. (2014) use a similar nonequilibrium phase transition model for protein accumulation in the endoplasmic reticulum to interpret experimental data on amyloyd-*β* clearance from the central nervous system.

## 9 Intrinsically disordered proteins

The relatively direct translation machinery of figure 1 is much more complicated in the case of intrinsically disordered proteins (IDP), or of more-or-less structured proteins having intrinsically disordered regions (IDR). Here, we adapt the approach of Wallace (2012d) to metabolic energy considerations.

The essential problem is that many proteins have no unique tertiary structure in isolation – or have regions without such structure – although they have distinct physiological function or functions in partnership with other chemical species. They lack a hydrophobic core, but lock with other molecules to engage in defined biological roles. That is, there is always some complex version of figure 1, implying the existence of an underlying code, and hence of some information channel.

Tompa et al. (2005) have observed that intrinsically disordered proteins provide unprecedented examples of protein signal moonlighting – multiple, often unrelated, functions of the same molecule – by eliciting both inhibiting and activating action on different partners, or even on the same partner. Figure 2, adapted from their paper, provides one schematic. The disordered protein can bind to more than one site on the partner molecule represented by a tilted square on the left of the figure. Binding to one site, as indicated by the shaded oval, creates an activated conformation, while binding to another site, the rectangle, results in an inhibited complex. Tompa et al. (2005) indicate, however, several different such possible mechanisms that are not mutually exclusive.

**Figure 2:**
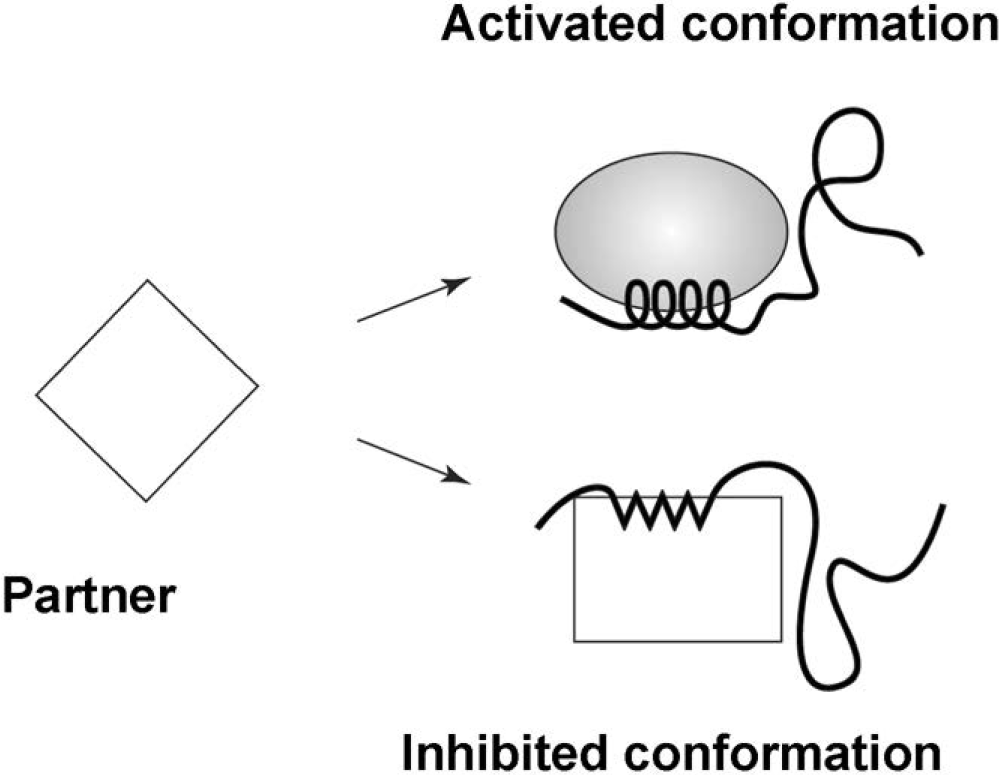
Adapted from Tompa et al. (2005). The partner, represented by the tilted square, can bind in two ways with the incoming IDP. The shaded oval represents activated, and the rectangle, inhibited species. The ‘choice’ between them is, in this model, to be made by an ‘information catalysis’ in which an incoming signal – itself requiring a minimum of metabolic free energy – shifts the lowest energy state between the two otherwise thermodynamically competitive conformations. This is one example of a vast spectrum of similar chemical ‘logic gates’.

We are interested in a single mode of such a switch, i.e., either the top or the bottom configuration. Assume it possible to extend nonrigid molecular group theory (Wallace 2012d) to the long, whip-like frond of an IDP/IDR anchored at both ends, via a sufficient number of semidirect and/or wreath products over an appropriate set of finite and/or compact groups. The dynamic mechanism of translation linkage with other chemical species is taken as parameterized by an index of ‘frond length’ *L* ∝ a binding energy index 𝓗 ≡ |𝓜| for some binding energy 𝓜 which might crudely be measured by the total number of amino acids in the IDP/IDR. At the very least, a potential energy barrier must be overcome for binding to take place. More generally, molecular species must be constructed and transported before binding, consuming metabolic free energies.

In general, the number of group elements can be expected to grow exponentially, as 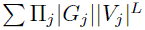, where |*G*_*k*_| and |*V*_*k*_| are the size, in an appropriate sense, of symmetry groups *G*_*k*_ and *V*_*k*_. Hence, for large *L* ∝ 𝓗, we are driven to a spontaneous symmetry shifting statistical mechanics approach on a Morse function, following the arguments of Pettini (2007) and Matsumoto (2002). Typically, many such Morse functions are possible, and it is possible to construct one using group representations.

Take an appropriate group representation by matrices and construct a ‘pseudo probability’ 𝒫 for nonrigid group element *ω* as

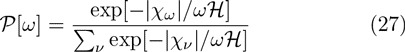

where *χ*_*ϕ*_ is the character of the group element *ϕ* in that representation, i.e., the trace of the matrix assigned to *ϕ*, and |…| is the norm of the character, a real number. For systems that include compact groups, the sum may be an appropriate generalized integral. The most direct assumption is that the representation is ‘faithful’, having as many matrices as there are group elements, but this may not be necessary.

The central idea is – again – that 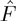 in the construct

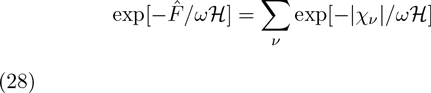

will be a Morse Function in the metabolic free energy measure 𝓗 to which we can apply Landau’s classic arguments on phase transition. Recall again the underlying idea, that, as the temperature of a physical system rises, more symmetries of the Hamiltonian become accessible, and this often takes place in a punctuated manner. As the temperature declines, these changes are characterized as ‘spontaneous symmetry breaking’. Here, we take the frond length *L* ∝ 𝓗 as the temperature index, and postulate punctuated changes in IDP/IDR function and reaction dynamics with its magnitude.

Recall that, given the powerful generalities of Morse Theory, virtually any good Morse Function will produce spontaneous symmetry shifts under these circumstances.

The observed ‘sloppiness’ of biological lock/key molecular reaction dynamics suggests that binding site symmetry may be greater than binding ligand symmetries: binding ligands may be expected to involve (dual, mirror) subgroups of the nonrigid group symmetries of the IDP/IDR frond.

A kind of ‘fuzzy lock theory’ emerges by supposing the ‘duality’ between a subgroup of the IDP/IDR and its binding site can be expressed as

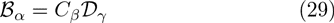

where *𝓑*_*α*_ is a subgroup (or set of subgroups) of the IDP/IDR nonrigid symmetry group, *𝓓*_*γ*_ a similar structure of the target molecule, and *C*_*β*_ is an appropriate inversion operation *or set of them* that represents static or dynamic matching of the fuzzy ‘key’ to the fuzzy ‘lock’.

If *C* is a single element, and *𝓑, 𝓓* fixed subgroups, then the matching would be classified as ‘static’. Increasing the number of possible elements in *C*, or permitting larger sets representing *𝓑* and *𝓓*, leads to progressively more ‘random’ structures in an increasingly dynamic configuration, as the system shifts within an ensemble of possible states, or, perhaps, even a superposition of them.

A more complete treatment probably requires a groupoid generalization of nonrigid molecule theory – extension to ‘partial’ symmetries like those of elaborate mosaic tilings, particularly for the target species. This approach has been highly successful in stereochemisty.

Again, molecular binding is not a free lunch. At the very least, potential energy barriers must be overcome. Such binding energy 𝓗 = |*𝓜*| must be provided by embedding metabolic free energy, and failure to deliver will trigger unwanted simplifications of physiologically essential IDP/IDR reactions.

## 10 A digression on free energy and regulation

More generally, the output of the logic gate in figure 2 can be viewed as constituting an information source – 0 for off, 1 for on – in which an external regulatory signal ‘chooses’ which configuration has the lower energy state. This can be formally expressed using the information theory chain rule (Cover and Thomas 2006). That is, information sources are often not independent, but are correlated, so that a joint information source can be defined having the properties

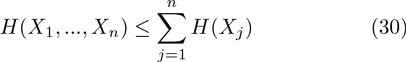

with equality only for isolated, independent information streams.

The chain rule has implications for free energy consumption in regulation and control processes. Again, Feynman (2000) describes how information and free energy have an inherent duality, defining information precisely as the free energy needed to erase a message. Information is a form of free energy and the construction and transmission of information within living things – the physical instantiation of information – consumes considerable free energy, with inevitable – and massive – losses via the second law of thermodynamics.

Suppose an intensity of available free energy is associated with each defined joint and individual information source having Shannon uncertainties *H*(*X*, *Y*), *H*(*X*), *H*(*Y*), e.g., rates 𝓗_*X,Y*_, 𝓗_*X*_, 𝓗_*Y*_.

Although information is a form of free energy, there is necessarily a massive entropic loss in its actual expression, so that the probability distribution of a source uncertainty *H* might again be written in Gibbs form as

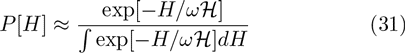

assuming *ω* is very small.

To first order, then,

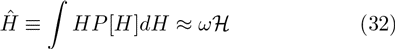

and, using equation (30),

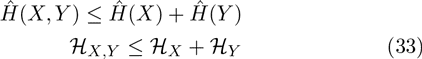

Thus, as a consequence of the information chain rule, allowing crosstalk consumes a lower rate of free energy than isolating information sources. That is, in general, it takes more free energy – higher total cost – to isolate a set of information sources than it does to allow them to engage in crosstalk.

Hence, at the free energy expense of supporting two information sources, – *X* and *Y* together – it is possible to catalyze a set of joint paths defined by their joint information source. In consequence, given a physiological system (or set of them) having an associated information source *H*(…), an external information source *Y* can catalyze the joint paths associated with the joint information source *H*(…,*Y*) so that a particular chosen reaction pathway – in a large sense – has the lowest relative free energy.

At the expense of larger global free information expenditure – maintaining two (or more) information sources with their often considerable entropic losses instead of one – the system can feed, in a sense, the generalized physiology of a Maxwell’s Demon, doing work so that regulatory signals can direct system response, thus locally reducing uncertainty at the expense of larger global entropy production.

That is, proper ongoing maintenance and operation of the IDP/IDR ‘switch’ implied by figure 2 – or any such logic gates – can require considerable metabolic free energy expenditure.

## 11 Glycan/lectin interaction

Glycan/lectin interaction at the surface of the cell, however, follows a different charateristic pattern. The actual ‘glycan/lectin logic switch’ has a markedly different dynamic from the IDP/IDR logic gate: no fuzzy-lock-and-key. Nonetheless, the MFE considerations of equations (30-33) still apply.

An example. The carbohydrate α-GalNAc interacts with the lectin biotinylated soybean agglutinin (SBA) in solution to form a sequence of increasingly complicated interlinked conformations at appropriate concentrations of reacting species. Dam et al. (2007) describe this ‘bind-and-slide’ process in terms of a change in topology, according to figure 3.

**Figure 3:**
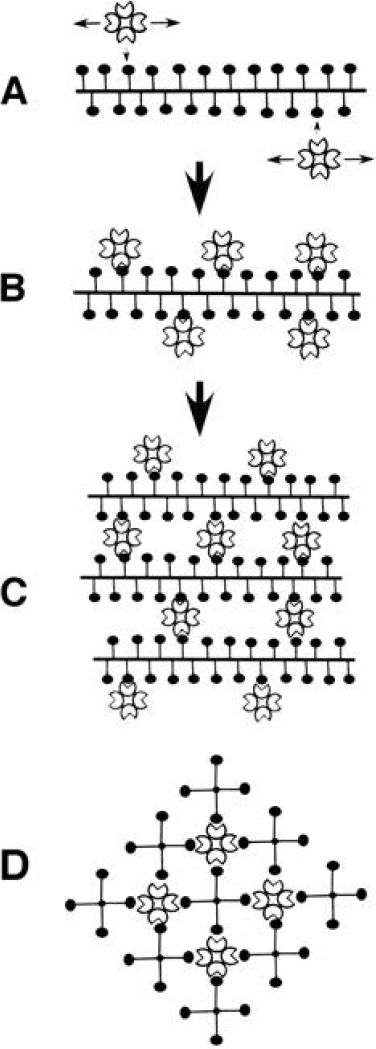
From Dam et al. (2007). (A) At first, lectin diffuses along and off the glycan kelp frond, until, (B), a sufficient number of sites are occupied. Then (C), the lectin-coated glycan fronds begin to cross bind and the reaction is saturated, and the gate thrown. (D) shows an end-on view of the complex in (C). A kind of spontaneous symmetry breaking with increasing lectin concentration is evident, with mode (A) far ‘freer’ than the locked-in state of modes (C) and (D). Details will vary with the particular glycan kelp frond and the impinging lectin species.

Initially, the lectin diffuses along (and off) the glycan kelp frond until a number of sites are occupied. Then the lectin-coated glycan fronds begin to cross bind, until the reaction saturates in a kind of inverse spontaneous symmetry breaking. Figure 3D shows an end-on view of the complex shown longitudinally in figure 3C.

Dam and Brewer (2008) generalize:

> The bind-and-slide model for lectins binding to multivalent glycosides, globular, and linear glycoproteins is distinct from the classical ‘lock and key’ model for ligand-receptor interactions. The bind and slide (internal diffusion) model allows a small fraction of bound lectin molecules to dynamically move from carbohydrate to carbohydrate epitope in globular and linear glycoproteins. This, in turn, can facilitate lectin-mediated cross-linking of such glycoproteins on the surface of cells… Such cross-linked receptors, in turn, trigger signal transduction mechanisms… Indeed, a large number of transmembrane receptors are found clustered… Thus the affinity and hence specificity of ligand-receptor interactions may be regulated by epitope and receptor clustering in many biological systems.

Under typical physiological circumstances, glycans form a literal kelp bed bound to cellular surfaces, and the essential topological ‘intensity parameter’ – the temperature analog – becomes area density of the fronds. See Dam and Brewer (2010) for details. Oyelaran et al. (2009), for example, conducted density-dependent fluorescence experiments, and it was possible to take the observed intensity of that fluorescence as an index of chemical information channel capacity and switch operation, since no information transmission indicates no reaction, producing no fluorescence.

While IDP/IDR switches appear directly amenable to a direct symmetry analysis, the construction and operation of ‘glycosynapses’ at the cell surface may require more study, both because of the apparently cognitive mechanisms producing the glycan kelp bed itself, and the diffusion-lock mechanism of the glycan/lectin switch.

Note that, from the perspective of Oyelaran et al., figure 3 could be reinterpreted as displaying a spontaneous symmetry breaking with increasing ‘kelp frond’ area concentration at a given lectin concentration: from the relatively free modes of 3A, to the suddenly locked-in ‘on’ state of 3C and 3D. Although this is examined by Wallace and Wallace (2013) using an IDP-like model, but it seems likely that a different approach may be needed. That is to say, the ‘code’ implied by figure 3 is necessarily different from the ‘codes’ implied by figures 1 and 2. Nevertheless, something much like equation (23) will dominate the stability of the basic switching mechanisms at the cell surface, since the transmission of information is, often critically, a matter of free energy availability and control signal strength.

## 12 The embedding symmetries of cognition

The arguments leading to equations (23), (25), and (30-33) are not restricted to face-to-face chemical interaction. Very many specific case histories, however, in the sense of Tomkins (1975), fall under the general rubric of a ‘metabolic code’:

> Complex regulation is characterized by two entities not operating in simple mechanisms: metabolic ‘symbols’ and their ‘domains’. The term ‘symbol’ refers to a specific intracellular effector molecule which accumulates when a cell is exposed to a particular environment… Metabolic symbols need bear no structural relationship to the molecules which promote their accumulation in a nutritional or metabolic crisis… Another important property of intracellular symbols is metabolic liability, which allows their concentrations to fluctuate quickly in response to environmental change…
>
> Since a particular environmental condition is correlated with a corresponding intracellular symbol, the relationship between the extra- and intracellular events may be considered as a ‘metabolic code’ in which a specific symbol represents a unique state of the environment…
>
> A second essential concept is complex regulation is that of the ‘domain’ of a symbol, defined as all the metabolic processes controlled by the symbol… [so that] the biochemical reactions included in the domain of a symbol are related by their biological effects rather than their chemical mechanisms…

This, however, is an argument perilously close to Atlan and Cohen’s (1998) characterization of cognition in biological systems. Under such circumstances, punctuated symmetry changes occur as groupoid shifts in cognitive function, rather than as simple amyloid-style shape change – an important observation that is worth some further comment.

Buried in the review by Motlagh et al. (2014) regarding the ensemble nature of biomolecular allostery is the striking statement that

> It is not obvious how the phenomenon of allostery can be understood and described in terms that can do equal justice to both highly structured and highly disordered systems.

Their figure 2 outlines an increasing dynamics of disorder or fluctuation in allostery, from rigid body motions, through side-chain dynamics, backbone dynamics, local unfolding, to intrinsically disordered systems.

Here, we have explored the metabolic dynamics of the often subtle symmetries of certain codes involved in the transmission of information within organisms, at several scales and levels of organization. There is, however, a more profound level of symmetry that dominates these structures and their dynamics. The various codes and blindingly complicated logic gates that are at the center of much current study instantiate cognitive mechanisms that, meshing together, constitute the living state. Very many cognitive processes can be characterized in terms of information sources having inherent symmetries, in a limited sense. The construction is curiously direct, and we follow the direction of Atlan and Cohen (1998), who first elucidated the ‘language of cognition’ for the immune system. They argue that the central feature of cognition is the comparison of a perceived signal with an internal, learned or inherited picture of the world, and then choice of one response from a much larger repertoire of possible responses. Thus cognitive pattern recognition-and-response proceeds by an algorithmic combination of an incoming external sensory signal with an internal ongoing activity, incorporating the internalized picture of the world, and triggering an appropriate action based on a decision that the pattern of sensory activity requires a response.

Incoming ‘sensory’ input – in a large sense – is, then, mixed in an unspecified but systematic manner with a pattern of internal ongoing activity to create a path of combined signals *x* = (*a*_0_, *a*_1_, …, *a*_*n*_,…). Each *a_k_* thus represents some functional composition of the internal and the external. An application of this perspective to a standard neural network is given in Wallace (2005).

This path is fed into a similarly unspecified ‘decision function’ *h,* generating an output *h*(*x*) that is an element of one of two disjoint sets *B_0_* and *B_1_* of possible system responses. Let *B*_*0*_ ≡ {*b*_0_, …, *b*_*k*_}, and *B*_*1*_ ≡ {*b*_*k*+1_, …, *b*_*m*_}.

Assume a graded response, supposing that if *h*(*x*) ∈ *B*_0_, the pattern is not recognized, and if *h*(*x*) ∈ *B*_1_, the pattern is recognized, and some action *b*_*j*_, *k* + 1 ≤ *j* ≤ *m* takes place.

Interest focuses on paths *x* triggering pattern recognition-and-response: given a fixed initial state *a*_0_, examine all possible subsequent paths *x* beginning with *a*_0_ and leading to the event *h*(*x*) ∈ *B*_1_. Thus *h*(*a*_0_, …, *a*_*j*_) ∈ *B*_0_ for all 0 ≤ *j* < *m*, but *h*(*a*_0_, …, *a*_*m*_) ∈ *B*_1_.

For each positive integer *n*, let *N*(*n*) be the number of high probability paths of length *n* that begin with some particular *a*_0_ and lead to the condition *h*(*x*) ∈ *B*_1_. Call such paths ‘meaningful’, assuming that *N*(*n*) will be considerably less than the number of all possible paths of length *n* leading from *a*_0_ to the condition *h*(*x*) ∈ *B*_1_.

Identification of the ‘alphabet’ of the states *a*_*j*_, *B*_*k*_ may depend on the proper system coarse graining, in the sense of symbolic dynamics (Beck and Schlogl, 1993).

Combining algorithm, the form of the function *h,* and the details of grammar and syntax, are all unspecified in this model. The assumption permitting inference on necessary conditions constrained by the asymptotic limit theorems of information theory is that the finite limit

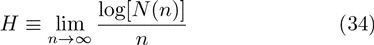

both exists and is independent of the path *x*. Recall that *N*(*n*) is the number of high probability paths of length *n*.

We define such a pattern recognition-and-response cognitive process as ‘ergodic’. Not all cognitive processes are likely to be ergodic, in this sense. Then *H*, if it exists in the limit *n* → ∞ (Khinchin 1957), is path dependent, although extension to nearly ergodic processes appears possible (Wallace 2005, pp. 31-32).

Now invoke the Shannon-McMillan Theorem. It is possible to define an adiabatically, piecewise stationary, ergodic information source **X** associated with stochastic variates *X*_*j*_ having joint and conditional probabilities *P*(*a*_0_, …, *a*_*n*_) and *P*(*a*_*n*_|*a*_0_, …, *a*_*n*−1_) such that appropriate joint and conditional Shannon uncertainties satisfy the classic relations of equation (7)

Indeed, this is one of the ‘master equations’ inherent to the nonequilibrium systems that generate information. As Khinchin (1957) emphasizes, nonergodic sources will have H-values that are path dependent, leading to ‘manifold’ structures (Wallace 2005, Section 3.1).

Such an information source will be called *dual* to the underlying ergodic cognitive process.

‘Adiabatic’ means that, when the information source is parameterized according to some appropriate scheme, within continuous pieces, changes in parameter values take place slowly enough so that the information source remains as close to stationary and ergodic as needed to make the fundamental limit theorems work. ‘Stationary’ means that probabilities do not change in time, and ‘ergodic’ (roughly) that cross-sectional means converge to long-time averages. Between pieces it is possible to invoke various kinds of phase change formalism, for example a biological form of standard renormalization theory (Wallace, 2005).

Recall that the Shannon uncertainties *H* (…) are cross-sectional law-of-large-numbers sums of the form −∑_*k*_ *P*_*k*_ log[*P*_*k*_], where the *P*_*k*_ constitute a probability distribution. See Cover and Thomas (2006) for the standard details.

We are not constrained to the Atlan-Cohen model of cognition, which involves representation, but can embrace as well more general insights. The basic idea is that a large class of cognitive phenomena – with or without representation – can be associated with a dual information source: cognition inevitably involves choice, choice reduces uncertainty, and this implies the existence of an information source, a language-analog.

The symmetries arise through an equivalence class algebra that can be constructed by choosing different origin points, *a*_0_, and defining the equivalence of two states, *a*_*m*_, *a*_*n*_, by the existence of a high probability meaningful path connecting them to the same origin point. Disjoint partition by equivalence class, analogous to orbit equivalence classes for dynamical systems, defines the vertices of a network of cognitive dual languages. Each vertex then represents a different information source dual to a cognitive process. This is not a representation of a physical network as such. It is, rather, an abstract set of ‘languages’ dual to the set of cognitive biological processes, and it is interactions across this set that will are of interest here.

Such a set of equivalence classes generates a groupoid, whose algebraic properties – an important extension of the idea of both a symmetry group and an equivalence class – are summarized in Wallace (2012a, c). For a groupoid, a product need not be defined globally. Again, see Brown (1987) and Weinstein (1996) for details, and the Mathematical Appendix for a summary.

Cognitive groupoids, in conjunction with the asymptotic limit theorems of communication and control theories, provide sufficient structure to impose significant ‘symmetry’ constraints on a vast range of biological subsystems.

Once the underlying groupoid nature of the cognitive processes that embrace ‘codes’, allostery, logic gates, and other constituent phenomena, is recognized, then it becomes possible to parse out the instantiating machinery according to dual information sources and their groupoid indices, defining another pseudoprobability in terms of available metabolic free energy as

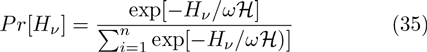

where *ν* represents the appropriate cognitive groupoid element. Then arguments like equations (24)-(26) can be imposed, or the dynamics explored using Onsager-like nonequilibrium thermodynamic formalism.

Define a ‘symmetry entropy’ based on the Morse Function *𝓕* arising from equation (35) over a set of structural parameters **Q** = [*Q*_1_, …, *Q*_*n*_] (that may include the rate of available metabolic free energy 𝓗 and other information source uncertainties) as the Legendre transform

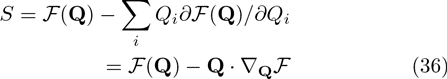

The dynamics of such a system will be driven, at least in first approximation, by Onsager-like nonequilibrium thermodynamics relations having the standard form (de Groot and Mazur, 1984):

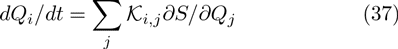

where the *𝓚*_*i*,*j*_ are appropriate empirical parameters and *t* is the time. A biological system involving the transmission of information may, or may not, have local time reversibility: in English, for example, the string ‘eht’ has a much lower probability than ‘the’. Without microreversibility, *𝓚*_*i*,*j*_ ≠ *𝓚*_*j*,*i*_.

Since, however, biological systems are quintessentially noisy, a more fitting approach is through a set of stochastic differential equations having the form

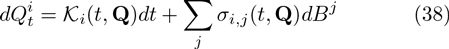

where the *𝓚*_*i*_ and *σ*_*i*,*j*_ are appropriate functions, and different kinds of ‘noise’ *dB*^*j*^ will have particular kinds of quadratic variation affecting dynamics (Protter, 1990).

Several properties become evident:

1. Setting the expectation of equation (38) equal to zero and solving for stationary points gives attractor states since the noise terms preclude unstable states.
2. This system may converge to limit cycle or pseudorandom ‘strange attractor’ dynamics.
3. What is converged to is not a simple state or limit cycle of states. Rather it is an equivalence class, or set of them, of highly dynamic modes coupled by mutual interaction through crosstalk and other interactions. Thus ‘stability’ in this structure represents particular patterns of ongoing dynamics rather than some identifiable static configuration. These are quasistationary nonequlibrium states.
4. Applying Ito’s chain rule for stochastic differential equations to the 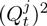 and taking expectations allows calculation of variances. These may depend very powerfully on a system’s defining structural constants, leading to significant instabilities depending on the magnitudes of the *Q*_*i*_, as in the Data Rate Theorem.
5. Following the arguments of Champagnat et al. (2006), this is very much a coevolutionary structure, where fundamental dynamics are determined by the feedback between ‘internal’ and ‘external’.

In particular, setting the expectation of equation (38) to zero generates an index theorem (Hazewinkel 2002) in the sense of Atiyah and Singer (1963), that is, an expression that relates analytic results – the solutions of the equations – to underlying topological structure, the eigenmodes of a complicated geometric operator whose groupoid spectrum represents ‘hidden symmetries’ of the overlying biocognitive dynamic that entrains the various constitutent allosteric and related molecular mechanisms.

## 13 Discusson and conclusions

For many ‘coded’ systems, some version of equations (23) and (30-33) will always emerge. Although equation (23) is built around rate distortion arguments for a Gaussian channel, the result extends to other channel architectures, since it is based on the convexity of the rate distortion function with the average distortion. Punctuated Data Rate Theorem instability failures, however, need not be restricted to the kind of symmetry collapse associated with amyloid formation.

Again, strictly speaking, biological codes, although they may be partly analyzed using information theoretic methods as Tlusty does, do not fall so easily within direct characterization by S/SM/RD/DR models. Such codes do not, in fact, invoke cognition, although, as argued above, they may become essential subcomponents within larger cognitive processes, reflecting the entraining symmetries of those processes.

Nonetheless, the argument leading to the Data Rate Theorem analog of equation (23) – and the generalization of Section 7 and equations (30-33) – place code stability within a recognizably similar framework, with MFE serving as a general ‘control signal’ roughly in the sense of Tomkins (1975), stabilizing efficient operation of complex biochemical coding and translation machinery, including – but not limited to – IDP/IDR mechanisms and the cell surface ‘Kelp bed’. Demand beyond available metabolic energy supply then expresses itself in punctuated destabilization, degradation, or pathological simplification, of the code/translation channel, possibly affecting gene expression, protein folding, IDP/IDR reactions, the operation of the glycan/lectin interface, and the more extensive mechanisms implied by Tomkins (1975).

Normal aging, or its acceleration by psychosocial or environmental stressors, must eventually interfere with routine code/translation operation – via deterioration in mitochondrial function and other possible mechanisms – triggering onset of many chronic diseases associated with senescence that involve failures of these essential biological processes.

By the development of Section 12, however, this may be a universal phenomenon, in which failing metabolic free energy degrades the full spectrum of biocognitive dynamics.

That is, the arguments of equations (15) and (23) – characterizing the role of metabolic free energy in the stability of biological control – and the ‘symmetry’ extension in Section 12, appear to be quite general, acting both at the kind of higher order cognitive functions implied by Maturana and Varela (1980) and Wallace (2014a, c), and at low level ‘basic’ code/translator scales, illuminating the role of mitochondrial deterioration in aging at all scales and levels of organization.

We have answered something of the question raised by Ge and Quan (2011) – and many others – regarding whether nonequilibrium phase transitions play an important role in living systems, and, perhaps, made some progress in resolving the allosteric paradox raised by Motlagh et al. (2014).

## 14 Mathematical Appendix

### The Data Rate Theorem

The Data-rate Theorem (DRT), a generalization of the classic Bode Integral Theorem for linear control systems, describes the stability of feedback control under data rate constraints (Nair et al., 2007). Given a noise-free data link between a discrete linear plant and its controller, unstable modes can be stabilized only if the feedback data rate ℐ is greater than the rate of ‘topological information’ generated by the unstable system. For the simplest incarnation, if the linear matrix equation of the plant is of the form *x*_*t*+1_ = **A***x*_*t*_ + …, where *x*_*t*_ is the n-dimensional state vector at time *t*, then the necessary condition for stabilizability is that

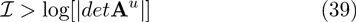

where *det* is the determinant and **A**^*u*^ is the decoupled unstable component of **A**, i.e., the part having eigenvalues ≥ 1. The determinant represents a generalized volume. Thus there is a critical positive data rate below which there does not exist any quantization and control scheme able to stabilize an unstable system (Nair et al. 2007).

The new theorem, and its variations, relate control theory to information theory and are as fundamental as the Shannon Coding and Source Coding Theorems, and the Rate Distortion Theorem for understanding complex cognitive machines and biological phenomena (Cover and Thomas, 2006). Very clearly, however, the DRT speaks to the centrality of the question whether nonequilibrium phase transitions play an important role in living systems.

### Groupoids

A groupoid, *G*, is defined by a base set *A* upon which some mapping – a morphism – can be defined. Note that not all possible pairs of states (*a_j_*, *a_k_*) in the base set A can be connected by such a morphism. Those that can define the groupoid element, a morphism *g* = (*a_j_*, *a_k_*) having the natural inverse *g*^−1^ = (*a_k_*, *a_j_*). Given such a pairing, it is possible to define ‘natural’ end-point maps *α*(*g*) = *a_j_*, *β*(*g*) = *a_k_* from the set of morphisms *G* into *A*, and a formally associative product in the groupoid *g_1_g_2_* provided *α*(*g*_1_*g*_2_) = α(*g*_i_),*β*(*g*_1_*g*_2_) = *β*(*g*_2_), and *β*(*g*_1_) = α(*g*_2_). Then, the product is defined, and associative, (*g*_1_*g*_2_)*g*_3_ = *g*_1_(*g*_2_*g*_3_). In addition, there are natural left and right identity elements λ*_g_*, *ρ_g_* such that λ*_g_g* = *g = gρ_g_*.

An orbit of the groupoid *G* over *A* is an equivalence class for the relation *a_j_* ∼ *Ga_k_* if and only if there is a groupoid element *g* with α(*g*) = *a_j_* and *β*(*g*) = *a_k_*. A groupoid is called transitive if it has just one orbit. The transitive groupoids are the building blocks of groupoids in that there is a natural decomposition of the base space of a general groupoid into orbits. Over each orbit there is a transitive groupoid, and the disjoint union of these transitive groupoids is the original groupoid. Conversely, the disjoint union of groupoids is itself a groupoid.

The isotropy group of *a* ∈ *X* consists of those *g* in *G* with α(*g*) = *a* = *β*(*g*). These groups prove fundamental to classifying groupoids.

If *G* is any groupoid over *A*, the map (α, *β*) : *G* → *A* × *A* is a morphism from *G* to the pair groupoid of *A*. The image of (*α*, *β*) is the orbit equivalence relation ∼ *G*, and the functional kernel is the union of the isotropy groups. If *f*: *X* → *Y* is a function, then the kernel of *f*, *ker*(*f*) = [(*x*_1_, *x*_2_) ∈ *X* × *X*: *f*(*x*_1_) = *f*(*x*_2_)] defines an equivalence relation.

Groupoids may have additional structure. For example, a groupoid *G* is a topological groupoid over a base space *X* if *G* and *X* are topological spaces and *α*, *β* and multiplication are continuous maps.

In essence, a groupoid is a category in which all morphisms have an inverse, here defined in terms of connection to a base point by a meaningful path of an information source dual to a cognitive process.

The morphism (*α*, *β*) suggests another way of looking at groupoids. A groupoid over A identifies not only which elements of A are equivalent to one another (isomorphic), but *it also parameterizes the different ways (isomorphisms) in which two elements can be equivalent,* i.e., in our context, all possible information sources dual to some cognitive process. Given the information theoretic characterization of cognition presented above, this produces a full modular cognitive network in a highly natural manner.

## 15 Acknowledgments

The author thanks Dr. D.N. Wallace for useful discussions.

